# Toxoplasma in a natural population of chimpanzees

**DOI:** 10.1101/2025.02.18.638806

**Authors:** Mathieu Chabry, Catherine Hobaiter, Klaus Zuberbühler, Jacob C. Koella

## Abstract

*Toxoplasma gondii* is a unicellular parasite that can infect most warm-blooded animals. In humans, it is linked to schizophrenia and other neurological disorders. It is also responsible for congenital toxoplasmosis, which can be lethal or cause severe neurological damage in the developing foetus. More subtle behavioural effects have been observed in various species, including humans, in particular increased risk-taking, which is thought to increase the parasite’s transmission to feline predators, the definitive hosts. We investigated the prevalence of *Toxoplasma gondii* in wild eastern chimpanzees (*Pan troglodytes schweinfurthii*) in the Budongo Forest, Uganda. We tested 45 individuals of two communities, one of which (Sonso) has a home range overlapping with human settlements. In this community, the prevalence was 22.7% (N=5 of 22), whereas in the neighbouring, but more isolated, Waibira community none of the 23 individuals tested were infected. The majority of infected Sonso individuals were adult males and/or high-ranking group members, attributes associated with increased risk-taking behaviour in chimpanzees and other animals. While further work is necessary to reveal the causal relation between infection and behaviour, our findings open new avenues of research for the neural impact of parasites on primate social behaviour.

## Introduction

*Toxoplasma gondii* is a protozoan parasite that infects about a third of the human population (Bigna et al. 2020), with prevalences ranging from 1% in Canada to over 60% in Ethiopia (Bisetegn et al. 2023). While the main threat is congenital toxoplasmosis during pregnancy (Furtado et al. 2011), which can lead to foetal death or blindness and neurological disorders later in life (Chaudhry et al. 2014), it has also been linked to diverse pathologies like epilepsy (Ngoungou et al. 2015) and schizophrenia (Contopoulos-Ioannidis et al. 2022), as well as to suicidal tendencies (Zerekidze et al. 2024). *Toxoplasma gondii* can also cause more subtle symptoms, including changes in behaviour (Flegr 2007), a possible consequence of the evolutionary pressure on the parasite’s life cycle to increase transmission. Although *T. gondii* can infect almost any warm-blooded animal as an intermediate host, its definitive hosts are members of the Felidae, which are infected when an intermediate host, typically a rodent, is eaten. To increase the probability that this happens, *T. gondii* appears to make mice and rats lose their natural fear of cat scents (Vyas et al. 2007a, 2007b) or to even become attracted to their urine (House et al. 2011). Similarly, infected hyena cubs are less risk-averse towards lions (Gering et al. 2021), wolves more likely to leave their natal groups (Meyer et al. 2022), and deer more likely to be killed by hunters (Nava et al. 2023). Loss of risk aversion has also been reported for infected primates, such as a decreased fear of leopard urine in captive apes (Poirotte et al. 2016) or, for humans, an enhanced risk of car accidents (Flegr et al. 2002; Yereli et al. 2006; Kocazeybek et al. 2009). Despite the importance of *T. gondii* for human health and its well-documented behavioural effects on laboratory animals, very little is known about the impact of this parasite in natural populations.

To address this, we investigated the prevalence of toxoplasmosis in two chimpanzee communities in the Budongo Central Forest Reserve, Uganda. One of them, the Sonso community, has a long history of cohabiting with humans, sharing part of its home range with a human settlement (a former sawmill) and regularly visiting the forest borders for crop-raiding. Such foraging behaviour is dangerous, can lead to fatal conflicts and is not practised by all group members (Tweheyo et al. 2005; Hockings and Humle 2009; McLennan and Hockings 2014; Anderson 2018). Our second study group, the Waibira community, has no known history of human contact, apart from a small team of researchers who have been studying them since the 2010s as well as infrequent contact with local people accessing the forest for resources.

We then investigated a possible association between *T. gondii* infection and socio-demographic variables to assess two possible relationships. First, high-ranked males are more likely to engage in risky behaviours than other group members, particularly in terms of territorial behaviour (Massaro et al. 2022), group hunting (Riedel et al. 2020), crop-raiding (Hockings et al. 2007) and anti-predator behaviour (Hockings et al. 2006). Some of these behaviours, in particular crop-raiding or entering villages, will bring them into close contact with situations in which they could be infected. The alternative hypothesis is that *T. gondii* infections occur more randomly but then secondarily affect the decision-making of infected individuals and so increase their chances of obtaining high social rank.

## Methods

We studied two groups of chimpanzees in the Budongo Central Forest Reserve in the western Rift Valley of Uganda, with a population of about 600 chimpanzees (Plumptre and Reynolds 1996). The two groups have been habituated to human presence and are followed by researchers and field assistants of the Budongo Conservation Field Station (www.budongo.org) on a daily basis. The Sonso group has been studied since 1990 and consisted of about 90 chimpanzees at the time of the study. Its home range borders on the forest edge in the south and overlaps with several human settlements. The Waibira group has been studied since 2011 and had about 120 group members at the time of the study.

Their territory is well within the main forest neighbouring the Sonso territory but located several kilometres from the nearest human settlements. All members of both communities are individually known, and for most it is possible to allocate at least ordinal ranks as high, medium or low.

We determined infection by *T. gondii* from urine samples collected while following focal animals between July and September 2024 throughout the day, usually from 7:30 to 16:30 local time (Sonso: 25 days; Waibira: 19 days). To estimate the prevalence in both communities, we aimed to collect two samples on different days from at least 20 adults per group, balanced for sex and including both alpha males. A focal animal was selected from the first subgroup (‘party’) encountered after entering the forest. We then determined the party composition and, if the alpha male was present, we selected him as the focal animal. The other focal animals were chosen to obtain an even adult male/female balance. Focal animals were followed at a distance of 7-10m on the ground, until we were able to collect a urine sample from leaves or the ground with a pipette. If the focal animal urinated from within a tree, we collected the falling urine with a plastic bag attached to a forked branch (following Crockford et al. 2013). The urine was stored in a vial, which was kept in a plastic bag until we returned to camp. If time permitted, we tried to collect urine from another individual on the same day.

Infection by *T. gondii* parasites was assessed by detecting the presence of IgG or IgM antibodies to the P30 antigen in the urine with the ID Screen Toxoplasmosis Indirect Multi-Species test (Innovative Diagnostics). The test kit is designed to work with blood, which made it necessary to first confirm that it can also reliably detect antibodies in urine. We therefore tested 11 adult humans, who all knew their *T. gondii* infection status (3 positive, 8 negative). The three positive individuals provided three urine samples, the 8 negative individuals at least 1 urine sample in 50 ml Falcon tubes. All samples were tested twice, mostly within 24 h of collection. The samples were centrifuged at 4000 g for 10 min in Amicon Ultra 10 filter tubes to increase the concentration of the antibodies and then tested with the test kit. All participants produced a *Toxoplasmas*-status that corresponded to their previously known status. In humans, *T. gondii* antibodies can be detected for years, possibly for life (Villard et al. 2016).

We tested chimpanzee urine with the same procedure, except that centrifugation was at 1250 g due to limitations in the field. All samples were tested on the same day that they were collected, and each sample was tested twice. For 29 individuals, we only managed to obtain one sample, as subjects temporarily left the core area of their territory. For the 16 individuals for which we had more than one sample, all samples gave the same result.

To test for possible associations between infection status and social variables (rank, sex, community membership), we used Fisher’s exact tests; and to test for an association between age and infection we used a two-sample t-test after having confirmed with Shapiro-Wilk tests that the distributions of age were normally distributed. Age classes were determined following Reynolds (2005).

## Results

We tested 45 individuals (22 from Sonso and 23 from Waibira, Table 1). Five of them had *T. gondii* antibodies in their urine. All of the positive individuals were adults (age range: 15 to 31 years) from the Sonso community (prevalence = 22.7%), whereas none of the Waibira individuals (age range: 9 to 44) tested positive (difference between communities: p = 0.021). There was no significant association between infection and adult age (t = 0.671, p = 0.506; Shapiro-Wilk normality test: positive individuals: W = 0.981, p = 0.940). Four of the five positive individuals were adult males (prevalence in Sonso adult males = 44.4%, Sonso adult females = 10.0%), but the difference between sexes was not significant (p = 0.382, two-tailed Fisher exact test). Four of the five positive individuals were high-ranking adults, including the alpha male (Table 1; prevalence high-ranking adults = 50.0%, medium-ranking = 16.6%, low-ranking = 0.0%), yielding a significant association between infection and social rank (p = 0.033, two-tailed Fisher exact test).

**Table 1.**
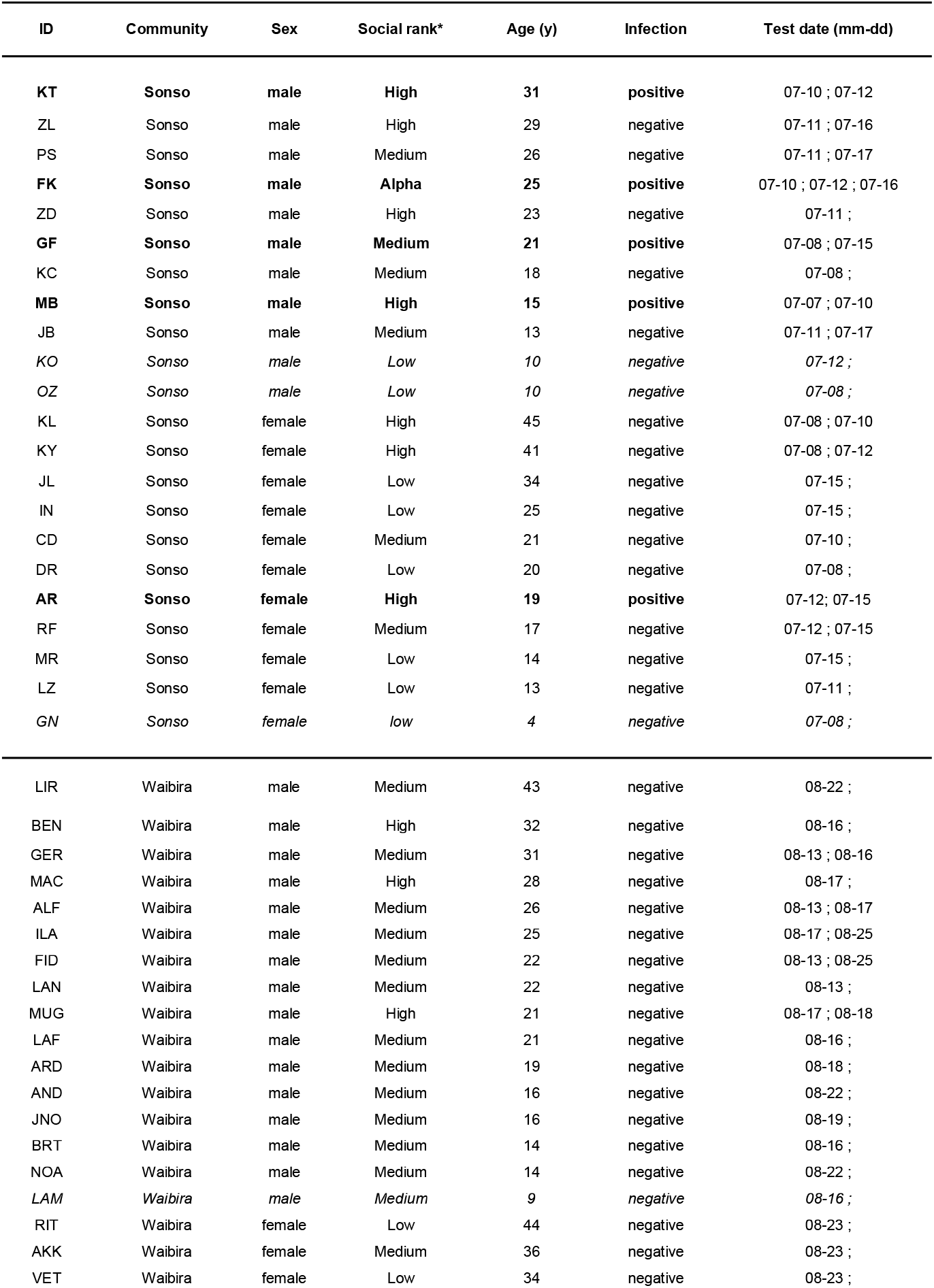
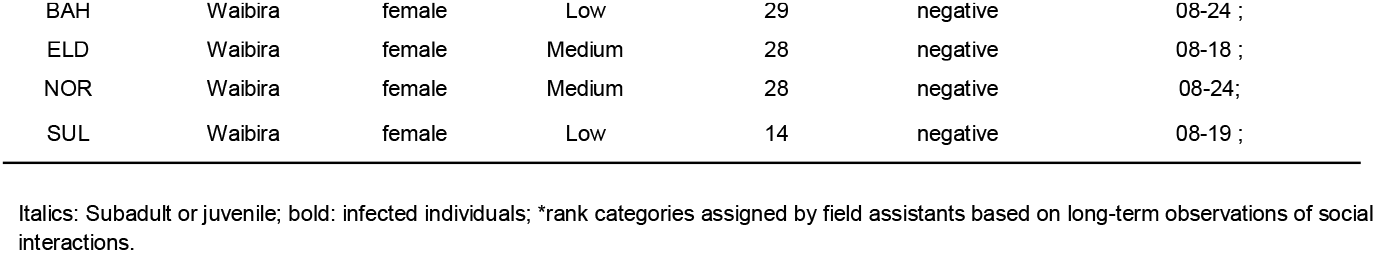
Demographic data and infection status of focal animals in two neighbouring chimpanzee communities.

| ID | Community | Sex | Social rank* | Age (y) | Infection | Test date (mm-dd) |
| --- | --- | --- | --- | --- | --- | --- |
| KT | Sonso | male | High | 31 | positive | 07-10 ; 07-12 |
| ZL | Sonso | male | High | 29 | negative | 07-11 ; 07-16 |
| PS | Sonso | male | Medium | 26 | negative | 07-11 ; 07-17 |
| FK | Sonso | male | Alpha | 25 | positive | 07-10 ; 07-12 ; 07-16 |
| ZD | Sonso | male | High | 23 | negative | 07-11 ; |
| GF | Sonso | male | Medium | 21 | positive | 07-08 ; 07-15 |
| KC | Sonso | male | Medium | 18 | negative | 07-08 ; |
| MB | Sonso | male | High | 15 | positive | 07-07 ; 07-10 |
| JB | Sonso | male | Medium | 13 | negative | 07-11 ; 07-17 |
| KO | Sonso | male | Low | 10 | negative | 07-12 ; |
| OZ | Sonso | male | Low | 10 | negative | 07-08 ; |
| KL | Sonso | female | High | 45 | negative | 07-08 ; 07-10 |
| KY | Sonso | female | High | 41 | negative | 07-08 ; 07-12 |
| JL | Sonso | female | Low | 34 | negative | 07-15 ; |
| IN | Sonso | female | Low | 25 | negative | 07-15 ; |
| CD | Sonso | female | Medium | 21 | negative | 07-10 ; |
| DR | Sonso | female | Low | 20 | negative | 07-08 ; |
| AR | Sonso | female | High | 19 | positive | 07-12 ; 07-15 |
| RF | Sonso | female | Medium | 17 | negative | 07-12 ; 07-15 |
| MR | Sonso | female | Low | 14 | negative | 07-15 ; |
| LZ | Sonso | female | Low | 13 | negative | 07-11 ; |
| GN | Sonso | female | low | 4 | negative | 07-08 ; |
| LIR | Waibira | male | Medium | 43 | negative | 08-22 ; |
| BEN | Waibira | male | High | 32 | negative | 08-16 ; |
| GER | Waibira | male | Medium | 31 | negative | 08-13 ; 08-16 |
| MAC | Waibira | male | High | 28 | negative | 08-17 ; |
| ALF | Waibira | male | Medium | 26 | negative | 08-13 ; 08-17 |
| ILA | Waibira | male | Medium | 25 | negative | 08-17 ; 08-25 |
| FID | Waibira | male | Medium | 22 | negative | 08-13 ; 08-25 |
| LAN | Waibira | male | Medium | 22 | negative | 08-13 ; |
| MUG | Waibira | male | High | 21 | negative | 08-17 ; 08-18 |
| LAF | Waibira | male | Medium | 21 | negative | 08-16 ; |
| ARD | Waibira | male | Medium | 19 | negative | 08-18 ; |
| AND | Waibira | male | Medium | 16 | negative | 08-22 ; |
| JNO | Waibira | male | Medium | 16 | negative | 08-19 ; |
| BRT | Waibira | male | Medium | 14 | negative | 08-16 ; |
| NOA | Waibira | male | Medium | 14 | negative | 08-22 ; |
| LAM | Waibira | male | Medium | 9 | negative | 08-16 ; |
| RIT | Waibira | female | Low | 44 | negative | 08-23 ; |
| AKK | Waibira | female | Medium | 36 | negative | 08-23 ; |
| VET | Waibira | female | Low | 34 | negative | 08-23 ; |
| BAH | Waibira | female | Low | 29 | negative | 08-24 ; |
| ELD | Waibira | female | Medium | 28 | negative | 08-18 ; |
| NOR | Waibira | female | Medium | 28 | negative | 08-24; |
| SUL | Waibira | female | Low | 14 | negative | 08-19 ; |

Italics: Subadult or juvenile; bold: infected individuals; \*rank categories assigned by field assistants based on long-term observations of social interactions.

## Discussion

We tested two neighbouring communities of eastern chimpanzees for the presence of *Toxoplasma gondii*. We found a surprisingly high prevalence of 22.7% in one of the two communities, Sonso, whose home range covers the Budongo Conservation Field Station, established in 1990 on the site of a former sawmill and extends to the forest edge. Sonso individuals seasonally visit human settlements and sugarcane plantations along the forest edge, which likely increases chances of contact with cat faeces containing resting stages of *T. gondii*. Furthermore, the field station is occasionally frequented by semi-feral cats, another possible source of contact and transmission.

All Waibira individuals tested negative. Although wild felids, i.e., golden cats and leopards, are present in other neighbouring forest reserves in Uganda, they have never been observed in the study area (C. Hobaiter, unpublished data). Moreover, the centre of the Waibira home range is about an hour’s walk from the Budongo Conservation Field Station and two hours from human villages. The Waibira community is forest-bound and surrounded by at least three other chimpanzee communities, including Sonso to the Southwest, and their only persistent contact with humans is through the research team at the Budongo Conservation Field Station. A similar inter-community difference has been reported for gastrointestinal parasites, with Sonso individuals displaying substantially higher *Ascaris* and cestode prevalence than Waibira (Yersin et al. 2017).

In the Sonso individuals we tested, age did not predict infection, but social rank did. One of the infected individuals of the Sonso group was the alpha male, and all others had a high social rank, apart from one middle-ranking male. This rank association could have emerged for two reasons. In line with the idea that *Toxoplasma* manipulates risk-aversion to enhance transmission, a *Toxoplasma* infection may indirectly impact the probability of obtaining high social status, most likely mediated in part by increased risk-taking behaviour (Holekamp and Strauss 2016; Funkhouser et al. 2018). Such a situation has been observed in wild wolves, where infected males were more likely to leave their natal groups and establish their own packs (Meyer et al. 2022). Other studies, however, found contradictory results regarding the relationship between infection and dominance (Gering et al. 2021). Alternatively, high rank may enhance the probability off *Toxoplasma* infection, for instance, if these individuals are likely to forage in or near human settlements where domestic cats are common, or if high-ranking individuals are more exploratory and therefore are more likely to contact cat faeces. Finally, although *Toxoplasma* can be sexually transmitted (Santana et al. 2013; Flegr et al. 2014; Hlaváčová et al. 2021), sexual transmission is unlikely to be a significant mechanism, as chimpanzee are highly promiscuous, which would lead to a high prevalence for all sexually active group members, irrespective of rank.

To distinguish between these possibilities, future research should include detailed behavioural analyses. In humans, there is a considerable literature with non-pathological differences between infected and non-infected individuals, including claims of personality change (Flegr 2007), impact on testosterone production (Hodková et al. 2007; Flegr 2010), changes in aggressive behaviour and impulsivity (Cook et al. 2015), mirroring some of the results also reported from rodents (Xiao et al. 2012; Golcu et al. 2014). As highlighted before, however, much of this literature is of a correlational nature (with a few exceptional studies in rodents; Berdoy et al. 2000; Vyas et al. 2007a, 2007b; Lamberton et al. 2008; Kannan et al. 2013; David et al. 2016; Boillat et al. 2020), and it remains largely unclear whether risk-prone individuals are more likely to get infected by *T. gondii* or the other way around.

### Conclusion and future directions

Here, we show that *T. gondii*, a parasite that can manipulate behaviour, can infect wild chimpanzees and highlight that further studies are needed to assess the impact of the parasite on the social structure and behaviour in groups, and their interaction with neighbouring groups. Of special importance is risk-taking behaviour, such as hunting, intra-group aggression, patrolling and inter-group aggression, foraging in the home range periphery and entering human habitats. The predictions are straightforward, with infected individuals more likely to engage in such behaviours than other group members, regardless of sex or community membership. It is possible that individuals who tested negative in this study will test positive in the future, which would allow us to draw more impactful conclusions about the causal relation between parasite and host, in the absence of experimentation. Also relevant is that chimpanzee communities differ dramatically in rates of violence, including lethal aggression, with isolated communities generally exhibiting lower rates of intra- and inter-group mortality than those with more extensive human contact (Wrangham et al. 2006). It is therefore possible that toxoplasmosis accentuates chimpanzees’ aggressive tendencies towards others (Wrangham et al. 2006; Wilson et al. 2014), suggesting that prevalence rates should be calculated for all long-term study sites. Furthermore, data on toxoplasmosis are not only important to understanding how this parasite affects the behaviour of the hosts, but may also serve as a basis to develop new tools in chimpanzee health monitoring. These could provide a valuable basis for designing novel conservation strategies and policies, particularly in light of the severe health implications of congenital toxoplasmosis in humans.

## Ethics

Permission to conduct the study and ethics approval was granted by the Uganda Wildlife Authorithy (permit number COD/96/05) and the Uganda National Council for Science and Technology (permit number NS789ES). Validation of the *Toxoplasma gondii* detection kit (ID Screen Toxoplasmosis Indirect Multi-Species test, Innovative Diagnostics) for urine was conducted with samples donated by people of known Toxoplasma status with their full informed consent. Samples were anonymised by names but infection status was known when testing. After testing all samples were destroyed in the laboratory.

## Data accessibility

Data can be accessed through this link: https://doi.org/10.5061/dryad.fqz612k36. AI has been used by M.C. to confirm the suitability of R codes and to improve their re-use.

## Author’s contributions

K.Z. and J.K. came up with the study, M.C. and J.K. designed the methods, M.C. conducted the experiments, M.C. wrote the original draft and M.C., K.Z., J.K. and C.H. wrote the final paper.

## Acknowledgements

We are grateful to the Royal Zoological Society of Scotland (Edinburgh Zoo) for providing long-term financial support for the Budongo Conservation Field Station (BCFS). At BCFS, we are grateful to David Eryenyu, Simon Peter Ogola, Suzan Lawino and Elijah Aliguma for logistical support. The study would not have been possible without the support and insights of Atayo Gideon and Monday Gideon, to whom we are especially grateful.

